# SignifiKANTE: Efficient *P*-value computation for gene regulatory networks

**DOI:** 10.1101/2025.11.13.688227

**Authors:** Fabian Woller, Paul Martini, Souptik Sen, David B. Blumenthal, Anne Hartebrodt

**Author notes:** These authors contributed equally to this work (Fabian Woller, Paul Martini). {, }.

## Abstract

Gene regulatory networks (GRNs) are graph-based representations of regulatory relationships between transcription factors and target genes. Various tools exist to infer GRNs from gene expression data, but since this task is computationally intensive, statistical significance estimates are often omitted. While permutation-based empirical *P*-value computation methods are relatively straightforward to implement, they are prohibitively expensive when applied to popular regression-based GRN inference methods and realistically sized datasets. To address this bottleneck, we developed SignifiKANTE. SignifiKANTE is based on the key insight that the background count distributions of groups of target genes may be highly similar, even if their expression vectors show distinct behavior. Relying on this insight, SignifiKANTE employs gene clustering based on the 1-Wasserstein distance to create a small, constant number of background distributions which enables the simultaneous computation of approximate empirical *P*-values for multiple target genes. This reduces runtime by orders of magnitudes (for some datasets, from several weeks to few hours), without compromising faithfulness of the obtained P-values. SignifiKANTE extends the popular GRN inference package Arboreto and is available as a Python package on GitHub (https://github.com/bionetslab/SignifiKANTE) and PyPI (https://pypi.org/project/signifikante/).

## 1 Introduction

Gene regulatory networks (GRNs) describe how transcription factors (TFs) [14] interact with target genes to control gene expression and enable the complex biological processes that support life. These networks provide key insights into physiological processes, like cell differentiation, disease progression, and response to external stimuli. Since experimentally determining regulatory links between TFs and their targets is expensive and such links can differ substantially between different conditions, tissues, and cell types, GRNs are often reconstructed from a high-throughput bulk or single-cell gene expression dataset *X* ∈ ℝ^*n×m*^ using computational methods (*n* denotes the number of samples or cells in the dataset, *m* denotes the number of genes). Various *in silico* GRN inference methods exist [20]. Typically, a TF-target gene edge is inferred between two genes if they are statistically associated across the samples (or cells in the case of single-cell data) in the dataset *X*. An edge-wise score indicating regulation strength is typically provided as well.

Here, we focus on one of the most common classes of GRN inference methods, which uses linear or non-linear regression to model the task of inferring a GRN from the gene expression data *X* [11,10,3]. These methods work as follows: Given a set of TFs *T* ⊆ {1, …, *m*} as candidate regulators, a regression model

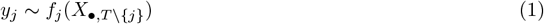

that predicts the expression profile *y*_*j*_ = *X*_•,*j*_ from the expression profiles of all TFs (except for *j* itself if *j* is a TF) is fit for each gene *j* ∈ {1, …, *m*.} Then, feature attribution methods are used that quantify the contributions of the TFs *t* ∈ *T* in the models *f*_*j*_. The obtained feature attribution scores are treated as weights *w*(*t, j*) of a potential regulatory link from the TF *t* to the target gene *j*, which are then aggregated across all targets *j* ∈ {1, …, *m*.} Finally, a GRN containing the links (*t, j*) with the largest weights is returned. To make the weights *w*(*t, j*) comparable across different target genes, the columns of *X* are typically normalized to unit variance before fitting the models *f*_*j*_ (e. g., using *z*-score normalization) [11]. Popular regression-based methods for GRN construction include GENIE3 [11] and GRNBoost2 [16]. GENIE3 employs random forest or extra tree regressors, which are replaced by gradient boosting with early stopping in GRNBoost2 to improve computational efficiency. Although regression-based methods such as GENIE3 and GRNBoost2 have performed very well in independent benchmarks [20], they share the common limitation that they rely on *ad hoc* cutoffs for the weights or the number of regulatory links to decide which edges (*t, j*) to include in the aggregated GRN. As a consequence, GRNs inferred by regression-based methods tend to include many false-positive regulatory links and have been shown to often lack robustness to random bias [12].

To address these issues and reduce the number of spurious interactions in inferred GRNs, statistical approaches to assess the significance of the edge weights *w*(*t, j*) are required. For GENIE3, such an approach has been developed as part of the DIANE dashboard for GRN inference and analysis [4]. In DIANE, empirical *P*-values are computed for weights *w*(*j, t*) through repeated shuffling of the target gene expression vectors *y*_*j*_, followed by random forest fitting with the permuted target vectors 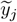. This approach generalizes to all regression-based GRN inference methods (see Algorithm A.1 in Appendix A) but suffers from severe scalability issues. It requires fitting *k* · *m* regression models, where *k* is the number of permutations determining the *P*-value resolution (typically *k* ≥ 1000). This is infeasible for GRNs inferred from realistically sized gene expression data with *m* ∈ [15000, 20000]. In DIANE, this problem is mitigated through two aggressive filtering steps prior to *P*-value computation: (i) restricting GRN inference to differentially expressed genes and (ii) applying a hard threshold on the weights *w*(*t, j*) to a achieve a user-specified (small) edge density. While these two filters make permutation-based *P*-value computation computationally feasible, they both involve additional hyperparameters without providing a clear rationale or data-driven strategy for their selection.

Beyond DIANE, statistical approaches for validating edges in regression-based GRNs have been proposed by Morgan et al. [18] and Wang et al. [23]. Morgan et al.’s strategy consists of repeated bootstrapping-based dataset resampling to improve stability and reduce false positives across a variety of regression-based GRN inference algorithms, including GENIE3 [11], LASSO [22,21], and CLR [6]. Wang et al. developed an approach employing perturbation-induced data coupled with bootstrapping and consensus-based link inclusion criteria to control the false discovery rate (FDR) in GRNs. However, both approaches have an even higher computational cost than the method implemented in DIANE and, in the respective publications, have been tested on data containing up to only 45 (Morgan et al. [18]) and 102 (Wang et al. [23]) genes.

With SignifiKANTE (from “signifikante Kante”, German for “significant edge”), we present the first algorithm for permutation-based assessment of edge significance in GRNs that overcomes these runtime limitations and scales to real-world gene expression data. SignifiKANTE makes critical use of the following key observation: In general, GRN inference methods pick up on statistical associations between two genes across the samples in the dataset. However, in the shuffled data, these statistical associations are removed. As a consequence, two genes with the same count distribution have identical null distributions, even if they are not statistically associated in the original expression data. Exploiting this observation, SignifiKANTE employs the following strategy to efficiently compute *P*-values for all edges in the originally inferred GRN: Instead of fitting one regression model for each of the *m* target genes for all *k* permutations, we first compute pairwise 1-Wasserstein distances between the column-normalized count distributions of all genes and use hierarchical clustering to create a constant number *ℓ* of null distribution prototypes. Based on these clusters, we create a proxy dataset containing representative genes for each cluster. This reduced dataset is then used to compute the null distribution by applying the user’s regression-based GRN inference method of choice. Since *ℓ* ≪ *m*, this substantially speeds-up the computation of *P*-values for all edges in a given GRN. SignifiKANTE is method-agnostic and can be used on top of every regression-based GRN inference approach.

To test SignifiKANTE’s approximation quality and computational efficiency, we compared against exact permutation-based *P*-values as implemented in DIANE, using 30 tissue gene expression datasets from GTEx [8]. Moreover, we compared GRNs with edges filtered based on SignifiKANTE’s *P*-values to vanilla GRNBoost2, relying on simulated data and comparisons of GRNs inferred from GTEx datasets to known TF-target gene links from CollecTRI [19], as well as their contextualization in the light of the BRENDA tissue ontology (BTO) [7]. SignifiKANTE is implemented in Python as an add-on for the widely used GRN inference package Arboreto [16], with out-of-the-box support for GENIE3, GRNBoost2, and LASSO-based GRN inference as implemented in GReNaDIne [21]. For users interested in using SignifiKANTE with other regression-based GRN inference methods, we provide a tutorial on how to easily integrate additional methods.

## 2 Results

### 2.1 Overview of SignifiKANTE

SignifiKANTE is based on the following key insight: If there are two target genes *j* and *j*′ whose columns 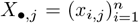 and 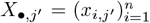 in the gene expression matrix *X* are identical up to sample permutation, then one of them is redundant under column-wise shuffling. When computing GRN edge weights upon column permutation to obtain DIANE-like empirical *P*-values, we can remove *j*′ from the set of targets and use edges targeting *j* as proxies for edges targeting *j*′. With this observation in mind, we call two gene expression vectors *X*_•,*j*_ and *X*_•,*j*_*′ equivalent up to sample permutation* if *sorted* (*X*_•,*j*_) = *sorted* (*X*_•,*j*_*′*), where *sorted* (·) denotes the sorting operation in non-decreasing order. Observation 1 characterizes gene expression vectors that are equivalent up to sample permutation in terms of the 1-Wasserstein distance.

#### Observation 1

*Two gene expression vectors X*_•,*j*_ *and X*_•,*j*_*′ are equivalent up to sample permutation if and only if W*_1_(*X*_•,*j*_, *X*_•,*j*_*′*) = 0, *where W*_1_(·, ·) *is the 1-Wasserstein or earth mover’s distance*.

#### Proof

Since *X*_•,*j*_ and *X*_•,*j*_*′* have the same length *n* and contain one-dimensional data, it holds that

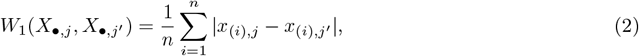

where the subscript (*i*) denotes the *i*^th^ order statistic. Since all summands in the right-hand side of Equation (2) are non-negative, *W*_1_(*X*_•,*j*_, *X*_•,*j*_*′*) = 0 holds if and only if |*x*_(*i*),*j*_ − *x*_(*i*),*j*_*′* | = 0 for all *i* = 1, …, *n*. By definition of the order statistics, this is the case if and only if *sorted* (*X*_•,*j*_) = *sorted* (*X*_•,*j*_*′*). □

SignifiKANTE (Figure 1A) exploits this observation as follows: We start by inferring a reference GRN from a column-normalized gene expression matrix and a list of TFs (candidate regulators) provided as input, using a regression-based GRN inference method of choice. Moreover, we group all target genes based on the 1-Wasserstein distance into a constant number of *ℓ* clusters. For a user-specified number *k* of permutations (*k* determines the *P*-value resolution, default: *k* = 1000), we then always draw one representative gene per cluster (through random selection or by computing the cluster medoid), permute its expression vector, and infer TF-target edges from the shuffled target gene vector. The resulting edge weights of the shuffled representative are extrapolated back onto all edges belonging to the respective cluster. We then count how often the edge weight inferred from the permuted data exceeds the corresponding reference edge weight, and turn this into an empirical *P*-value per edge. We thus obtain *P*-values for all edges in the reference GRN, which can optionally be adjusted for multiple testing using Benjamini-Hochberg (BH) correction.

**Fig. 1.**
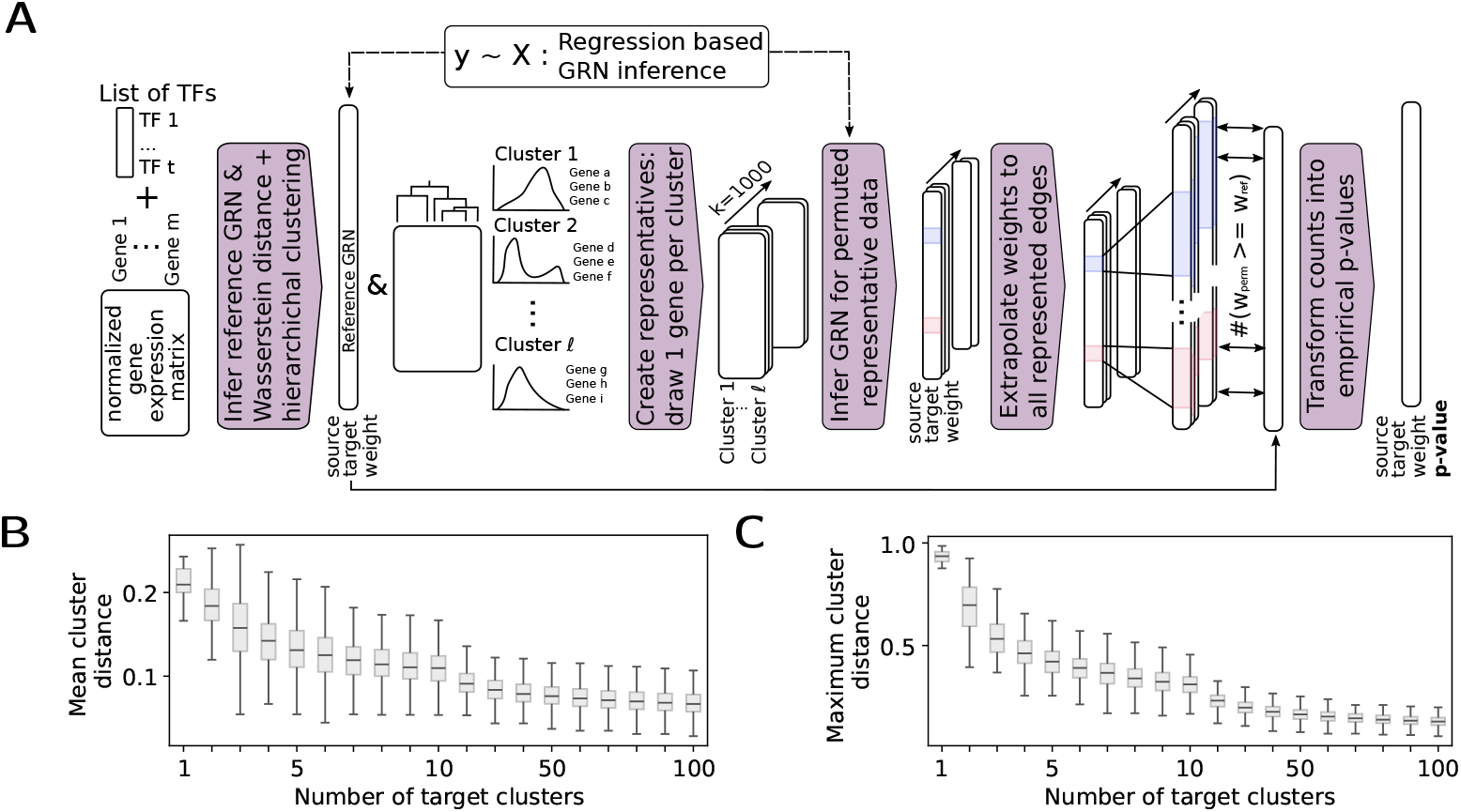
Method overview and assessment of distributional gene expression diversity in GTEx data. **(A)** As input, SignifiKANTE takes a gene expression matrix with columns scaled to unit variance (e. g., using *z*-score normalization) in combination with a regression-based GRN inference method of choice that infers a set of weighted TF-target edges from the data (reference GRN). Target genes are clustered using 1-Wasserstein distance-based hierarchical clustering, the regression-based GRN inference method is run *k* times with shuffled gene expression data for one representative per cluster, and the resulting edge weights between TFs and cluster representatives are used to compute approximate empirical *P*-values for all edges in the reference GRN. **(B, C)** Distributions of mean and maximum pairwise intra-cluster 1-Wasserstein distances resulting from 1-Wasserstein distance-based hierarchical clustering of gene expression profiles for 30 tissue gene expression datasets obtained from GTEx.

When analyzing the clustering structure resulting from the Wasserstein-based hierarchical clustering, we observed a common pattern across *z*-score normalized gene expression data of all 30 GTEx tissues (Figure 1B, C). Both mean and maximum pairwise intra-cluster distances show a pronounced decrease for one up to 10 target gene clusters, with only moderate improvements from 20 up to 100 clusters. This observation shows that distributional diversity of column-normalized gene expression vectors is relatively small, suggesting that already 10 to 100 clusters should yield accurate approximate *P*-values.

For all of the experimental results presented in the following sections, we ran SignifiKANTE with *k* = 1000 permutation and GRNBoost2 as the core GRN inference method. We selected GRNBoost2 as default because it is one of the most popular regression-based GRN inference methods and has been shown to yield a good tradeoff between runtime and accuracy in previous benchmarks [20]. However, since SignifiKANTE allows for easy integration of any regression-based GRN inference method, we also carried out analyses on a subset of the test datasets using GReNaDIne’s LASSO regression and GENIE3 as the core GRN inference method. The corresponding results are shown in Figure B.2 in Appendix B. As input list of candidate regulators, we used the widely used list of human TFs from the Stein Aerts Lab [1].

### 2.2 SignifiKANTE reduces runtime by orders of magnitude and yields highly accurate approximate *P*-values

First, we assessed accuracy of SignifiKANTE’s approximate *P*-values in comparison to exact empirical *P*-values computed with the DIANE-like approach explained in the introduction and in Algorithm A.1 in Appendix A. For this, we used 30 healthy tissue gene expression datasets from GTEx (see Table B.2 in Appendix B for an overview of the datasets), ran GRNBoost2 to compute reference GRNs for all of them, and computed exact and approximate *P*-values, using *k* = 1000 permutations for both exact and approximate *P*-values and a varying number 1 ≤ *ℓ* ≤ 100 of target gene clusters for the approximate *P*-values. Treating edges with exact empirical *P*-values below 0.05 (with and without BH correction) as groundtruth, we then computed F1 scores for edges with significant approximate *P*-values, using the same significance thresholds.

Figure 2 shows the results with random selection of cluster representatives (using medoids as cluster representatives performed slightly worse, see Figure B.1 in Appendix B), aggregated over tissue datasets with similar numbers of samples (*n <* 500, 500 ≤ *n* ≤ 1000, and *n >* 1000). SignifiKANTE was orders of magnitude faster than the exact DIANE-like *P*-value computation, reducing runtimes from days or weeks to hours. Even when using *ℓ* = 100 target gene clusters, SignifiKANTE runtimes fall below three hours on average on large datasets and below one hour on smaller ones (Figure 2A). When using *ℓ* = 10 target gene clusters, average runtimes of little more than one hour can be achieved also for the large datasets. Depending on the dataset sizes and the number of target gene clusters, these runtimes correspond to average speedup factors between 80 on small datasets and 270 on larger ones (Figure 2B), resulting in an average 80-to 400-fold reduction in estimated CO_2_ emission (Figure B.1 in Appendix B).

**Fig. 2.**
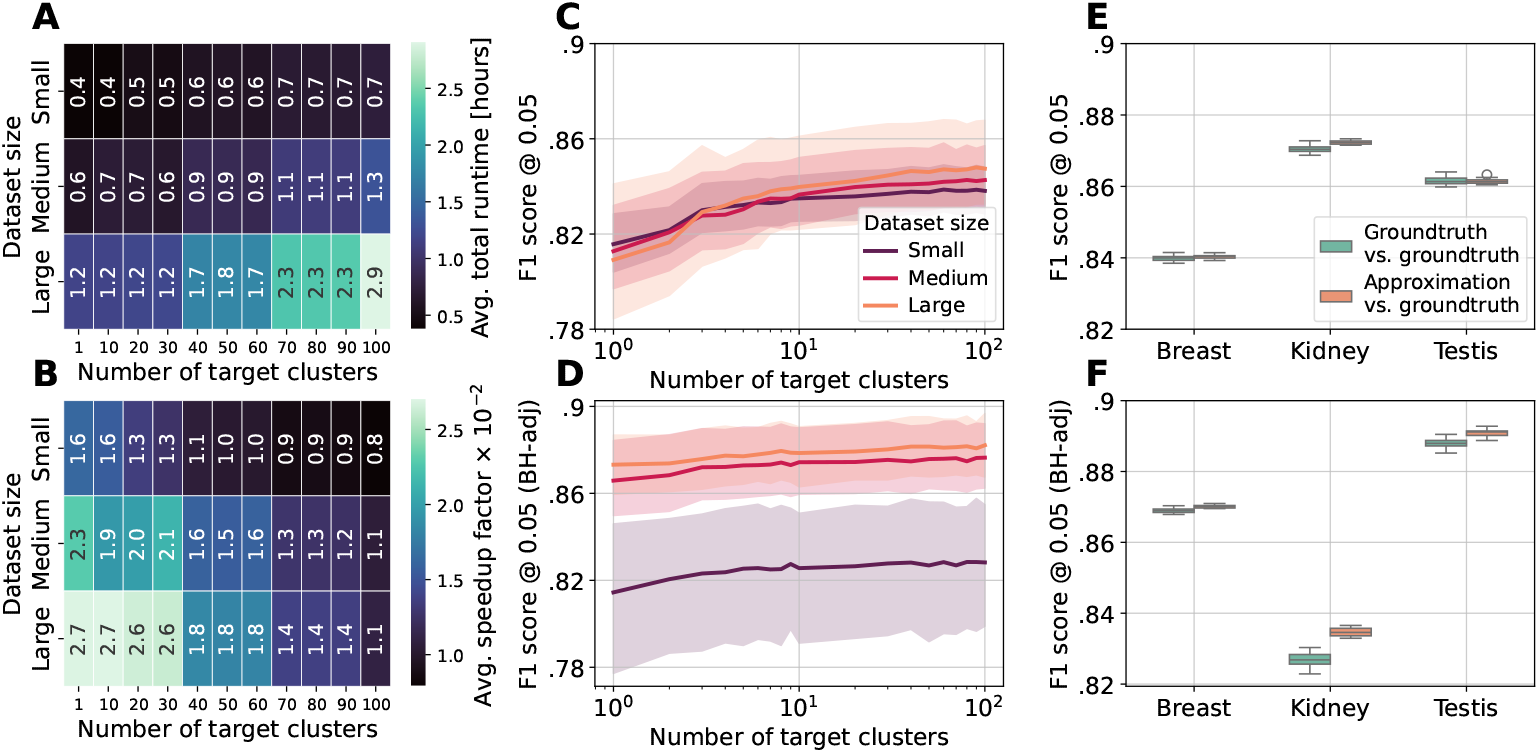
Comparison of SignifiKANTE to exact empirical *P*-value computation for GRNs as implemented in DIANE [4]. **(A)** Mean SignifiKANTE runtimes across small (*n <* 500), medium (500 ≤ *n* ≤ 1000), and large (*n >* 1000) GTEx datasets for varying *ℓ*. **(B)** Corresponding mean speed-up factors in comparison to exact *P*-value computation. **(C, D)** F1 scores for varying *ℓ* obtained through comparison of edges in GRNs with significant exact and approximate *P*-values at 0.05 significance level, both without (C) and with (D) BH correction to adjust for multiple testing. **(E, F)** F1 scores measuring approximation quality of SignifiKANTE’s *P*-values against ten groundtruth *P*-values and of ten groundtruth *P*-value runs against themselves for significance level 0.05, both without (E) and with (F) BH correction.

Regarding the accuracy of the approximate *P*-values (Figure 2C, D), we observe that, already for *ℓ* = 10 target gene clusters, F1 scores always lie above a minimal value of around 0.78, with a further improvement for increasing *ℓ*. Moreover, F1 scores tend to be higher on tissues with larger sample sizes, especially when applying BH correction (Figure 2D). To put these F1 scores in the context of the intrinsic stochasticity of permutation-based empirical *P*-values, we ran ten independent groundtruth calculations for a subset of three tissues (since already one groundtruth computation took days to weeks, doing this for all 30 tissues was computationally infeasible). We then analyzed pairwise F1 scores among all combinations of the ten groundtruths by pretending one of the two to be the “actual groundtruth” and the other one to be the “approximation”. Figures 2E, F present pairwise F1-scores of the ten groundtruths against themselves (“Groundtruth vs. groundtruth”) and against SignifiKANTE’s approximate *P*-values, using *ℓ* = 100 target gene clusters (“Approximate vs. groundtruth”). Our results demonstrate that, both with and without BH correction, the F1 scores achieved by SignifiKANTE coincide with the F1 scores of the pairwise groundtruth comparisons. This shows that the approximation quality of SignifiKANTE’s *P*-values is already optimal up to variation in the computation of the exact permutation-based *P*-values.

### 2.3 SignifiKANTE improves precision of inferred GRNs

Next, we asked to which extent SignifiKANTE improves upon the underlying GRN inference method GRN-Boost2. For this, we simulated gene expression datasets in such a way that a given groundtruth GRN has a strong effect on the genes’ expression profiles, used vanilla GRNBoost2, and GRNBoost2 with SignifiKANTE (with and without BH correction) to infer GRNs from the simulated datasets, and then assessed the inferred GRNs’ precisions in comparison to the groundtruth GRNs underlying the simulation. Figures 3A and B show the obtained precisions for the top *κ* % of edges with the largest weights in the different networks for varying *κ*, as well as pairwise Jaccard indices between the GRNBoost2 edge sets and the edge sets obtained with SignifiKANTE. The underlying groundtruth GRN consists of five TFs and 50 targets per TF, while precision values for two further groundtruth configurations are shown in Figure B.3 in Appendix B. For edges with large weights, the edge sets for the filtered and unfiltered networks are close to identical and hence yield very similar precision values. This indicates that edges with a high weight are generally statistically stable and have a significant *P*-value, which is an expected result. However, in the lower ranking edges, SignifiKANTE improves the precision on the simulated data, by removing statistically weak edges — especially when used in combination with BH correction. These results suggest that SignifiKANTE’s significance filtering effectively separates relevant from non-relevant low-weight TF-target gene links. Moreover, we observed that the relation between edge weight percentiles and *P*-values is often shifted between sets of outgoing edges incident with different TFs (Figure 3C). This implies that, with a purely weight-based GRN filtering strategy that does not make use of SignifiKANTE’s *P*-values, one either retains many non-relevant edges for TFs where the relation between weight percentiles and *P*-values is left-shifted (turquoise TF1 in Figure 3C) or looses relevant ones for TFs where this relation is shifted to the right (orange TF1 in Figure 3C).

**Fig. 3.**
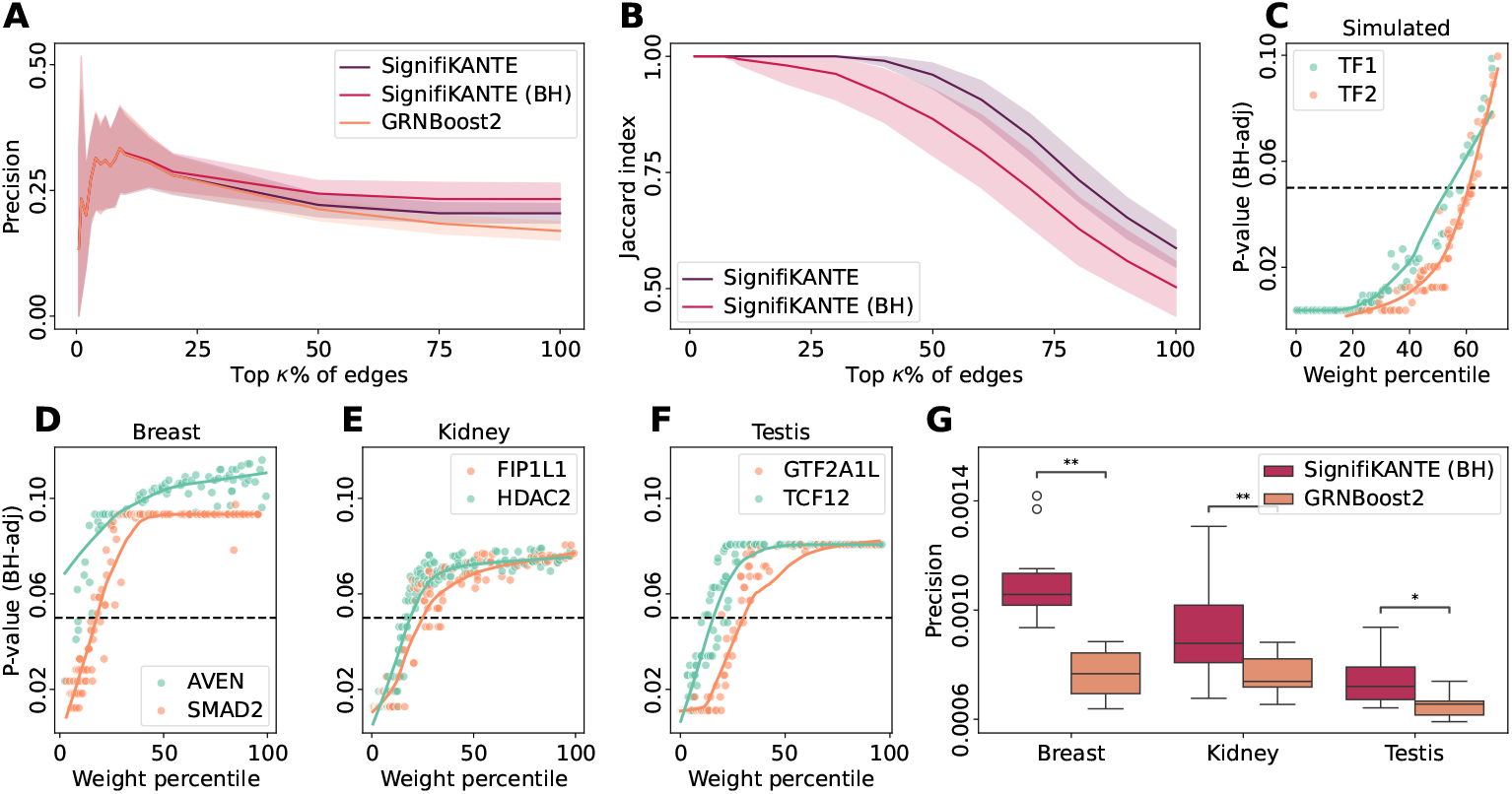
Effect of significance filtering with SignifiKANTE on GRNs inferred with GRNBoost2 from simulated and real-world data. **(A)** Precision of top *κ* % edges (sorted by weights) inferred from simulated expression data against groundtruth GRN used for simulation, using all GRNBoost2 edges, all significant edges of SignifiKANTE (significance cutoff at *α* = 0.05, *ℓ* = 10 target gene clusters), and all significant edges of SignifiKANTE with BH correction (FDR control at *q* = 0.05, *ℓ* = 10). The groundtruth GRN includes five TFs and at most 50 targets per TF. **(B)** Pairwise Jaccard indices between GRNBoost2 and SignfiKANTE edge sets for the edge cutoffs from (A). **(C–F)** Edge weight percentiles (low percentiles correspond to large weights) and BH-adjusted SignifiKANTE *P*-values for TF-target gene links of exemplary pairs of TFs on simulated data (C, FDR control at *q* = 0.05, *ℓ* = 10), and GTEx data for breast, kidney, and testis (D–F, FDR control at *q* = 0.05, *ℓ* = 100). Regression lines are fitted using locally weighted scatterplot smoothing. **(G)** Distributions of precision values between all GRNBoost2 edges inferred from breast, kidney, and testis GTEx data and only significant edges computed by SignifiKANTE (BH correction, and FDR control at *q* = 0.05, *ℓ* = 100). Known TF-target interactions from the CollecTRI database [19] were treated as groundtruth. Using different random seeds, we ran GRNBoost2 followed by SignifiKANTE ten times on the same data, and then assessed statistical significance via the Wilcoxon signed-rank test for matched samples.

In addition to our tests on simulated data, we evaluated the ability of SignifiKANTE to filter out irrelevant edges from the given reference GRNBoost2 GRN on a subset of three GTEx tissues. For this, we first filtered the respective reference GRNs for the most expressive regulatory interactions by subsetting it to edges targeting the 1000 most highly expressed target genes and then compared the GRNs of vanilla GRNBoost2 to GRNs filtered by SignifiKANTE (with BH correction, since this yielded the best results on simulated data). As for simulated data, we observe TF-specific shifts in the relation between edge weight percentiles and SignifiKANTE’s *P*-values (Figure 3D–F). This shows that, also for real-world data, the commonly used GRN filtering strategy to consider only the top *κ* % of edges with the largest weights either leads to a systematic inclusion of spurious links for some TFs (turquoise TFs in Figure 3D–F) or to a systematic exclusion of relevant links for other TFs (orange TFs in Figure 3D–F). Furthermore, on the same set of GTEx tissues, we compared the enrichment of known TF-target interactions from the CollecTRI database [19] within the complete GRNBoost2 GRNs and SignifiKANTE’s filtered FDR-controlled GRNs (Figure 3G). For all three tissues, the precision values computed when treating CollecTRI as groundtruth increase slightly but significantly compared to the vanilla GRNBoost2 GRNs. We also computed recall values corresponding to the precision values shown in Figure 3A, G (Figure B.3 in Appendix B). However, since SignifiKANTE computes subnetworks of the GRNs inferred by GRNBoost2, we know *a priori* that recall is higher for GRNBoost2 than for the filtered networks, which is why precision is the more relevant metric for our study.

### 2.4 SignifiKANTE improves representation of the BRENDA tissue ontology

Finally, we analyzed how well SignifiKANTE’s FDR-controlled GRNs inferred for the 30 GTEx tissue datasets reflect known (dis-)similarites between tissues. For this, we computed GRN-based tissue distances for all 30 GTEx tissues and compared them to distances based on the BTO [7], which we calculated by mapping the GTEx tissues to corresponding BTO terms (Table B.1 in Appendix B) and then computing pairwise shortest path distances on the ontology graph. Pairwise GRN-based tissue distances were calculated as Jaccard distances of the edge sets, using the edge sets inferred by vanilla GRNBoost2 and the significant edges resulting from, respectively, exact DIANE-like *P*-values and approximate *P*-values computed with SignifiKANTE with 100 target gene clusters, both coupled with BH correction with FDR control at *q* = 0.05.

Figure 4 shows how well the respective GRN-based tissue distances correlate with the BTO-based distances. Our analysis revealed that FDR-controlled GRNs from SignifiKANTE yield a higher Spearman rank correlation coefficient against BTO-based tissue distances than both DIANE-like FDR control and vanilla GRNBoost2 (Figure 4A). An even more pronounced gap can be observed when filtering the reference GRN to only contain the 1000 most highly expressed genes per tissue (Figure 4C). The corresponding *P*-values for all Spearman correlation coefficients are significant (Figure 4B, D). We can thus conclude that significance-based edge filtering with SignifiKANTE leads to GRNs that better reflect onotology-based tissue similarities than unfiltered GRNs inferred by vanilla GRNBoost2.

**Fig. 4.**
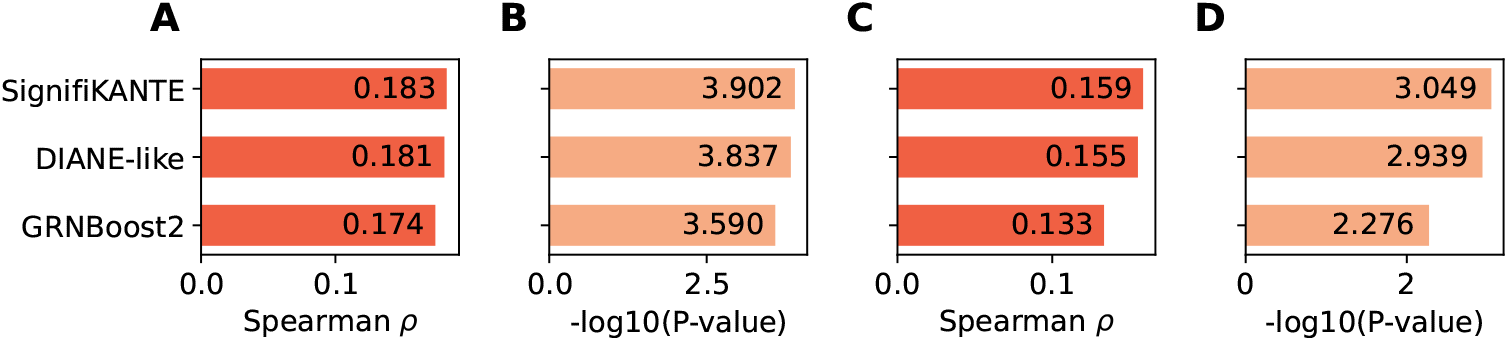
Comparison of edge similarity between GRNs inferred for different GTEx tissues with the BRENDA tissue ontology. **(A)** Spearman correlations of edge-wise Jaccard distances between GRNs and BTO-based shortest path distances across all 30 GTEx tissues, using GRNs containing, respectively, all GRNBoost2 edges and all significant edges for BH-corrected (FDR control at *q* = 0.05) exact DIANE-like *P*-values and approximate *P*-values from SignifiKANTE (*ℓ* = 100). **(B)** *P*-values corresponding to Spearman correlation coefficients shown in (A). **(C)** Spearman correlations of BTO-based shortest path distances and edge-wise Jaccard distances between GRNs, with GRNs filtered to the 1000 most highly expressed target genes. **(D)** *P*-values corresponding to Spearman correlation coefficients shown in (C).

## 3 Discussion

In this article, we presented SignifiKANTE — the first tool that allows to assess edge significance in GRNs inferred from realistically sized gene expression data without requiring *ad hoc* filtering steps to decrease the computational burden. For this, SignifiKANTE computes approximate empirical *P*-values using clustering based on the 1-Wasserstein distance. In terms of quality, the approximate *P*-values match exact empirical *P*-values for GRN edges suggested in a previous study [4], but can be computed in minutes to few hours instead of days to weeks. SignifiKANTE can be used on top of any regression-based GRN inference method; as default, we suggest the widely used method GRNBoost2 which was also used for most of our analyses. In comparison to GRNs inferred with vanilla GRNBoost2, SignifiKANTE’s GRNs show improved quality, especially in the tail of the TF-target gene links inferred by GRNBoost2 (for the edges with the highest weights, differences are small because most highly weighted edges are statistically significant). While the quality improvement is rather small in terms of magnitude, it is consistent across various metrics and datasets (precision in comparison to groundtruth and literature-curated TF-target gene links for simulated and real-world data, consistency with tissue ontology) and mostly statistically significant. In view of these results, we are convinced that SignifiKANTE is a valuable contribution to the network biology field and will help to make GRN inference from gene expression data more robust and reliable.

Nonetheless, our study has several limitations. Firstly, SignifiKANTE is tied to regression-based GRN inference and cannot be used on top of other approaches such as the mutual information-based method ARACNe-AP [13]. Secondly, our evaluation of SignifiKANTE when used with core GRN inference methods other than GRNBoost2 remains relatively limited — mainly due to the huge computational cost of computing the exact empirical *P*-values required for the evaluation. Finally, we did not test SignifiKANTE on single-cell gene expression data, e. g., when used to robustify the core GRNBoost2-based GRN inference step within the widely used single-cell GRN inference tool SCENIC [3]. Since single-cell gene expression data are much more noisy than bulk data, we anticipate that significance filtering with SignifiKANTE may have an even stronger effect when used for single-cell data. In future work, we will explore this possibility.

## 4 Methods

### 4.1 Pseudocode of SignifiKANTE

A high-level algorithmic description of SignifiKANTE is shown in Algorithm 1. As input, SignifiKANTE requires a gene expression matrix *X*, a set of column indices *T* denoting TFs (candidate regulators), a regression-based GRN inference method `GRN_inference()` together with the matching aggregation operator ⊕ to aggregate TF-target gene links for different target genes, and two technical hyperparameters *ℓ, k* ∈ ℕ. We start by normalizing the columns of the gene expression matrix to unit variance (Line 1, e. g., using *z*-score normalization), and then infer the GRN (*E, w*) for our data using `GRN_inference()` and ⊕ (Line 2). Next, we compute a matrix *D* ∈ ℝ^*m×m*^ containing 1-Wasserstein distances for all pairs of columns in *X* (Line 3), and then use *D* to compute a gene clustering via agglomerative hierarchical clustering (Line 4). The number of gene clusters *ℓ* is a hyperparameter that has to be set by the user. Larger *ℓ* result in a more fine-grained representation of the original gene expression space but increase SignifiKANTE’s runtime.

#### Algorithm 1

SignifiKANTE

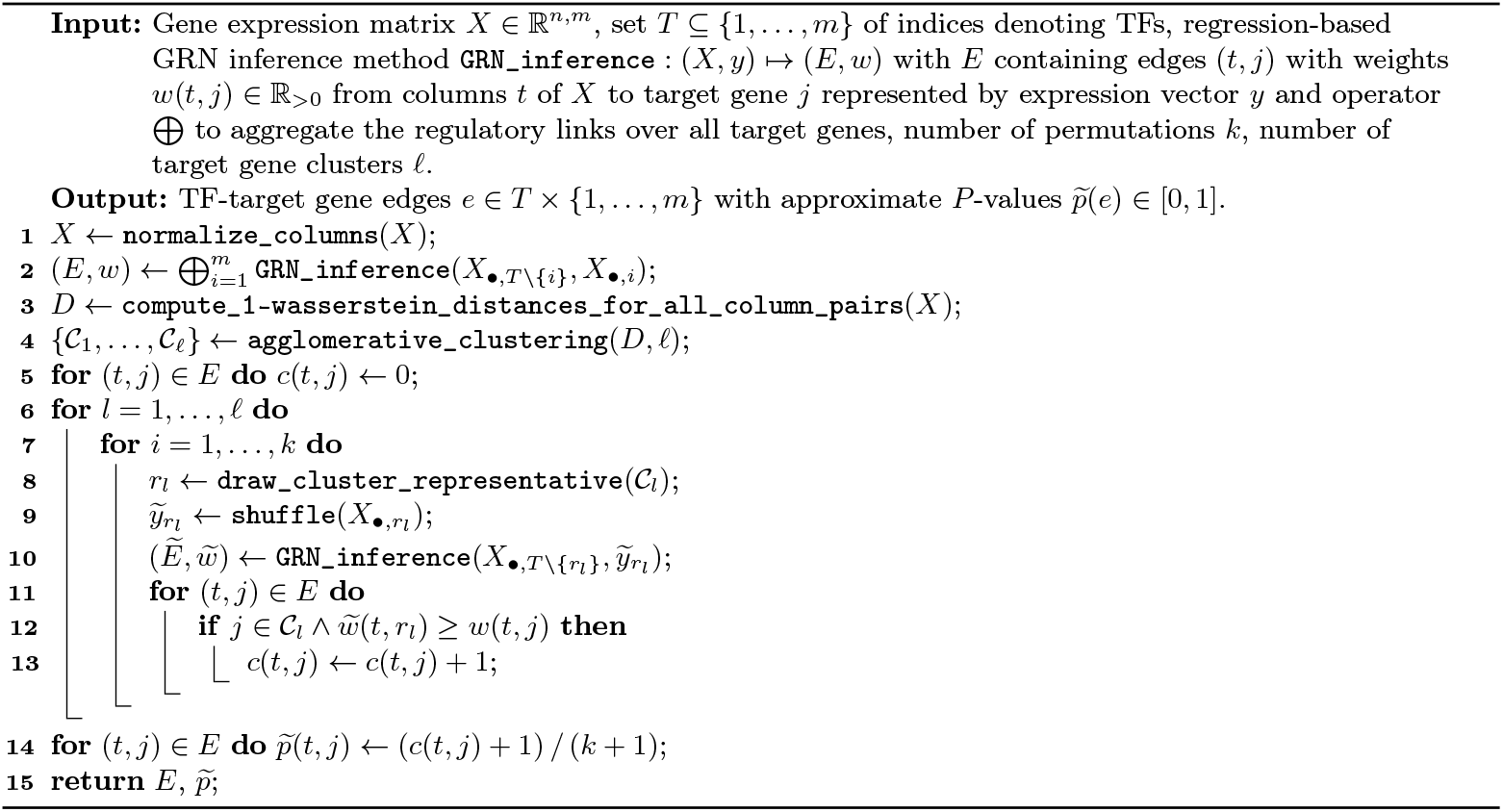

We then compute approximate empirical *P*-values for all edges (*t, j*) ∈ *E* using a sped-up version of the approach presented in DIANE, parametrized with the user-specified number of permutations *k*. For each edge (*t, j*) ∈ *E*, we maintain a zero-initialized integer *c*(*t, j*) (Line 5) that counts how often weights obtained with `GRN_inference` upon permutation of the target gene expression vector exceed the weight *w*(*t, j*) obtained with the real data. However, instead of computing a full ramdomized GRN for each of the *k* rounds of permutations as in DIANE, we only compute background weights for edges towards one representative *r*_*l*_ per gene cluster 𝒞_*l*_ in each iteration (Lines 9, 10). For drawing *r*_*l*_ (Line 8), we implemented two strategies: either using the cluster medoid (computed based on the 1-Wasserstein distance) or drawing a random cluster representative in each iteration. For the update of the edge counts *c*(*t, j*), the target gene *j* of a given edge (*t, j*) ∈ *E* from the reference GRN *G* is mapped to its corresponding cluster representatives *r*_*l*_ and the edge count of *c*(*t, j*) is increased if the weight 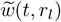 of the representative edge in the GRN obtained upon permutation of column *r*_*l*_ is higher than the we ight in the reference GRN *G* (Lines 11–13).

Once *k* randomized GRNs have been computed for representatives of all *ℓ* target gene clusters, SignifiKANTE computes approximate empirical *P*-values for all edges (*t, j*) ∈ *E* in the GRN inferred from the original data (Lines 14, 15). These *P*-values can optionally be adjusted for multiple testing (with the number of tests set to |*E* |) using BH correction, which is natively supported in the SignifiKANTE Python package. Note, however, that FDR cutoffs on the adjusted *P*-values should be chosen in light of the resolution 1*/*(*k* +1) of the empirical *P*-values. For instance, with SignifKANTE’s default *k* = 1000, the smallest possible non-adjusted empirical *P*-value is just below 0.001, and in preliminary experiments we thus obtained mostly empty result sets when controlling the FDR at *q* ≤ 0.01. This motivated us to select *q* = 0.05 as default.

### 4.2 Implementation

For computing the matrix 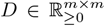 of pairwise 1-Wasserstein distances in Line 3 of Algorithm 1, we implemented a more efficient equivalent of SciPy’s `wasserstein_distance()` function, which allows us to handle large expression matrices with several thousand gene columns. To accomplish this, we relied on parallelization and code compilation to C using the Python just-in-time compiler Numba [2]. This reduced the runtime required to generate *D* to the order of minutes. Agglomerative hierarchical clustering based on *D* (Line 4 in Algorithm 1) was implemented using sklearn’s `cluster.AgglomerativeClustering` class to obtain *ℓ* target gene clusters. Note that by setting *ℓ* = *m*, the user can also employ SignifiKANTE to compute exacte empirical *P*-values as in DIANE. Since GRNBoost2 in its Arboreto implementation is one of the most popular GRN inference tools, we decided to implement SignifiKANTE as an add-on to Arboreto. Arboreto uses Dask for parallelizing the main loop over the target genes in regression-based GRN inference (that is, the regression models *y*_*j*_ ~ *f*_*j*_(*X*_•,*T \*{*j*}_) are fit in independent Dask processes). For SignifiKANTE, we changed the logic of the parallelization to avoid the repeated costly creation and shutdown of Dask processes in each permutation. Instead of making a full call to Arboreto’s GRN inference function for each permutation, we parallelize the loop over the target gene clusters (Line 6 in Algorithm 1) and, within each Dask process, compute the empirical *P*-values for the partial GRN containing edges towards target genes from the respective cluster. SignifiKANTE provides out-of-the-box support for the two GRN inference methods GRNBoost2 and GENIE3 implemented in Arboreto, as well as for GReNaDIne’s LASSO-based approach. The latter implementation also serves as a template which showcases how users can easily extend SignifiKANTE with any regression-based GRN method beyond the ones implemented in Arboreto.

### 4.3 Real-world and simulated gene expression data

We used both real-world and simulated gene expression data for our study. Real-world bulk RNA-sequencing data for 30 healthy tissues was obtained in transcripts per million (TPM) from GTEx and was used with minimal processing (column-wise *z*-score normalization, genes expressed in fewer than 10% of samples filtered out, numbers of samples and genes after preprocessing are shown in Table B.2 in Appendix B). The simulated data underlying our comparison of GRNBoost2 with and without edge filtering based on SignifiKANTE’s *P*-values was generated with the scMultiSim package [15], which allows to simulate gene expression data from an underlying groundtruth GRN. We used subnetworks of the CollecTRI GRNs as a base for our simulation, randomly subsampled to *s* TFs and min {*t, deg* (*i*) } unique target genes per randomly sampled TF *i* (*deg* (*i*) is the outdegree of the TF *i* in CollecTRI), with (*s, t*) ∈ { (5, 50), (10, 40), (20, 30) }. Using scMultisim, and setting a strong induced effect for these subnetworks, we then simulated 10 gene expression datasets with *n* = 500 cells for each (*s, t*) configuration, containing *m* ∈ {82, 159, 286} genes on average.

### 4.4 Evaluation metrics for comparison against exact DIANE-like *P*-values

For each edge *e* ∈ *E* in the reference GRN inferred from a GTEx dataset *X*, we computed exact *P*-values *p*(*e*) with the DIANE-like approach (Algorithm A.1 in Appendix A) and approximate *P*-values 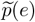 with SignifiKANTE. Due to the immense computational cost of the exact *P*-value computation task and a job time limit of 24 hours on the HPC systems provided by the Erlangen National High Performance Computing Center (NHR@FAU), we chunked the list of target genes into several batches for each tissue. For this, we set up a Slurm array job, which could then iterate over all tissues’ target gene batches and compute exact *P*-values on the respective chunk in under 24 hours, using Dask parallelization on 30 CPU cores. Like this, we were able to compute exact *P*-values for all 30 GTEx datasets on our HPC environment. Runtimes were recorded using Python’s `time` module. To compute the runtimes per tissue for the exact *P*-value computation that exceeded our 24 hours job time limit, we summed up the runtimes of the corresponding target gene batches. CO_2_ emission of exact and approximate *P*-value computation was estimated based on the `OfflineEmissionsTracker` module from the CodeCarbon Python package [5].

Given the exact and approximate *P*-values, we computed corresponding multiple testing-adjusted *P*- values *p*_BH_(*e*) and 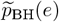, using BH correction. Then, we constructed four edge sets *E*^⋆^ = {*e* ∈ *E* | *p*(*e*) *<* 0.05}, 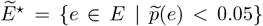, 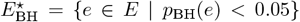, and 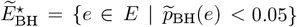, and computed F1 scores treating *E*^⋆^ as groundtruth for 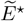 and 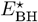 as groundtruth for 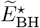. For the “Groundtruth vs. groundtruth” and “Approximation vs. groundtruth” (Figure 2E, F), we computed 10 variants 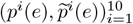 of the exact and approximate empirical *P*-values using different random seeds for the reference GRN inference, leading to edges sets 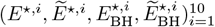. Then, we computed distributions of groundtruth-vs.-groundtruth F1 scores by comparing 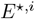 against 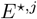 and 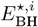 against 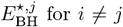 for *I* ≠ *j*, and approximation-vs.-groundtruth F1 scores by comparing 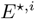 against 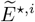 and 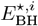 against 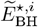.

### 4.5 Evaluation metrics for comparison against vanilla GRNBoost2

For the comparison on the simulated data, we constructed edges sets *E*^*κ*^ as the sets of top *κ* % of edges (sorted by edge weights) of the edge set *E* inferred by vanilla GRNBoost2, as well as edge sets 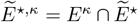 and 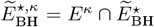 (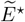 and 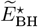 defined as above). For 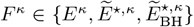, we then computed precision and recall at *κ* as precision(*F* ^*κ*^) = |*F* ^*κ*^ ∩ *E*_GT_|*/*|*F* ^*κ*^| and recall(*F* ^*κ*^) = |*F* ^*κ*^ ∩ *E*_GT_|*/*|*E*_GT_|, where *E*_GT_ is the groundtruth network from the data simulation. We also computed pairwise Jaccard indices 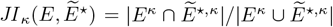 and 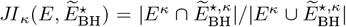. Note that 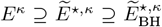 always holds, which implies recall 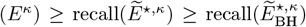. For the precision, there is no such *a priori* known monotonous relationship, which makes it the more relevant metric. For the GTEx data, we computed precision(*F*) = |*F* ∩ *E*_CollecTRI_|*/*|*F* | and recall(*F*) = |*F* ∩ *E*_CollecTRI_|*/*|*E*_CollecTRI_| for 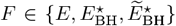, where *E* and 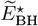 are defined as before, 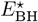 is the corresponding FDR-controlled GRN based on exact DIANE-like *P*-values, and *E*_CollecTRI_ is a set of curated TF-target gene edges obtained from CollecTRI, which we used as reference. Due to the incompleteness and non-specificity of CollecTRI, *E*_CollecTRI_ cannot be seen as a real gold-standard. However, it can be used to evaluate if the biological signal is improved.

To assess the correlation of pairwise edge-based Jaccard distances of GRNs with ontology-based tissue distances, we first conducted a manual mapping of the 30 GTEx tissue types onto BTO terms (see Table B.1 in Appendix B). The graph representing the BTO was then downloaded from the project website in the `.obo` file format and read into a NetworkX [9] graph object. Using NetworkX, we then computed pairwise shortest path distances between the mapped BTO terms in order to obtain our ontology-based distance matrix of tissues, against which we then compared our GRN-based distance matrices.

### 4.6 Computing environment used for benchmarking

All benchmarks have been conducted on the HPC resources provided by NHR@FAU. More specifically, we ran our benchmarks on the CPU-based compute cluster “Woody”, which is equipped with one Intel Xeon E3-1240 v5 (“Skylake”) CPU, one Intel Xeon E3-1240 v6 (“Kaby Lake”) CPU, and two Intel Xeon Gold 6326 (“Ice Lake”) CPUs. In all our benchmarks, we used a total of 30 cores for the parallelization with Dask.

## 5 Code and data availability

SingifiKANTE’s source code is available at https://github.com/bionetslab/SignifiKANTE, with an easy-to-install packaged version under https://pypi.org/project/signifikante, and scripts to reproduce results under https://github.com/bionetslab/SignifiKANTE_Results. The FDR controlled GRNs on all thirty GTEx tissues are publicly available at https://doi.org/10.5281/zenodo.17581025. The list of human TFs is taken from https://resources.aertslab.org/cistarget/tf_lists/, and GTEx data from https://www.gtexportal.org/home/downloads/adult-gtex/bulk_tissue_expression. CollecTRI was accessed using the decoupler package [17], and the BTO from https://www.brenda-enzymes.org/ontology.php?ontology_id=3.

## Acknowledgments

F. W. and D. B. B. were funded by the German Research Foundation (DFG, 516188180). D. B. B. and A. H. were funded by the German Federal Ministry of Research, Technology and Space (BMFTR, 031L0309A). P. M. and D. B. B. were funded by the Bavarian Center for Cancer Resaerch (BZKF, TLG/24/02/Krau). The authors gratefully acknowledge the scientific support and HPC resources provided by the Erlangen National High Performance Computing Center (NHR@FAU) of the Friedrich-Alexander-Universität Erlangen-Nürnberg (FAU). The hardware is funded by the German Research Foundation (DFG).

## Disclosure of Interests

All authors declare no competing interests.

## Notes

### Competing Interest Statement

The authors have declared no competing interest.

