## Appendices for "SignifiKANTE: Efficient *P*-value computation for gene regulatory networks"


Fabian Woller[1,†], Paul Martini[1,†], Souptik Sen[1,2], David B. Blumenthal[1], and Anne Hartebrodt[1]

[1] Biomedical Network Science Lab, Friedrich-Alexander-Universität Erlangen-Nürnberg, Erlangen, Germany
{fabian.woller,paul.martini,david.b.blumenthal,anne.hartebrodt}@fau.de

[2] Peter L. Reichertz Institut für Medizinische Informatik, Medizinische Hochschule Hannover, Germany



#### A Exact empirical $P$-value computation for regression-based GRN inference

Algorithm A.1 shows how to compute empirical $P$-values for GRNs inferred from gene expression data using regression-based approaches. The algorithm generalizes the approach that is implemented for GENIE3 [4] in the DIANE dashboard for GRN inference and analysis [1] to arbitrary regression-based GRN inference methods. Note that the two filtering steps used in DIANE prior to $P$-value computation (restricting GRN inference to differentially expressed genes and applying a hard threshold on the weights $w(t, j)$ to a achieve a small edge density) are not reflected in Algorithm A.1.

**Algorithm A.1:** DIANE-like empirical $P$-value computation for regression-based GRN inference

**Input:** Gene expression matrix $X \in \mathbb{R}^{n,m}$, set $T \subseteq \{1, \ldots, m\}$ of indices denoting TFs, regression-based GRN inference method `GRN_inference` : $(X, y) \mapsto (E, w)$ with $E$ containing edges $(t, j)$ with weights $w(t, j) \in \mathbb{R}_{>0}$ from columns $t$ of $X$ to target gene $j$ represented by expression vector $y$ and operator $\bigoplus$ to aggregate the regulatory links over all target genes, number of permutations $k$.

**Output:** TF-target gene edges $e \in T \times \{1, \ldots, m\}$ with empirical $P$-values $p(e) \in [0, 1]$.

$X \leftarrow$ `normalize_columns`$(X)$;
$(E, w) \leftarrow \bigoplus_{j=1}^{m}$ `GRN_inference`$(X_{\bullet,T\setminus\{j\}}, X_{\bullet,j})$;
**for** $(t, j) \in E$ **do** $c(t, j) \leftarrow 0$;
**for** $j = 1, \ldots, m$ **do**
  **for** $i = 1, \ldots, k$ **do**
    $\widetilde{y}_j \leftarrow$ `shuffle`$(X_{\bullet,j})$;
    $(\widetilde{E}, \widetilde{w}) \leftarrow$ `GRN_inference`$(X_{\bullet,T\setminus\{j\}}, \widetilde{y}_j)$;
    **for** $(t, j) \in E$ **do**
      **if** $\widetilde{w}(t, j) \geq w(t, j)$ **then**
        $c(t, j) \leftarrow c(t, j) + 1$;
**for** $(t, j) \in E$ **do** $p(t, j) \leftarrow (c(t, j) + 1) \,/\, (k + 1)$;
**return** $E$, $p$;

† These authors contributed equally to this work.

### B Supplementary figures and tables

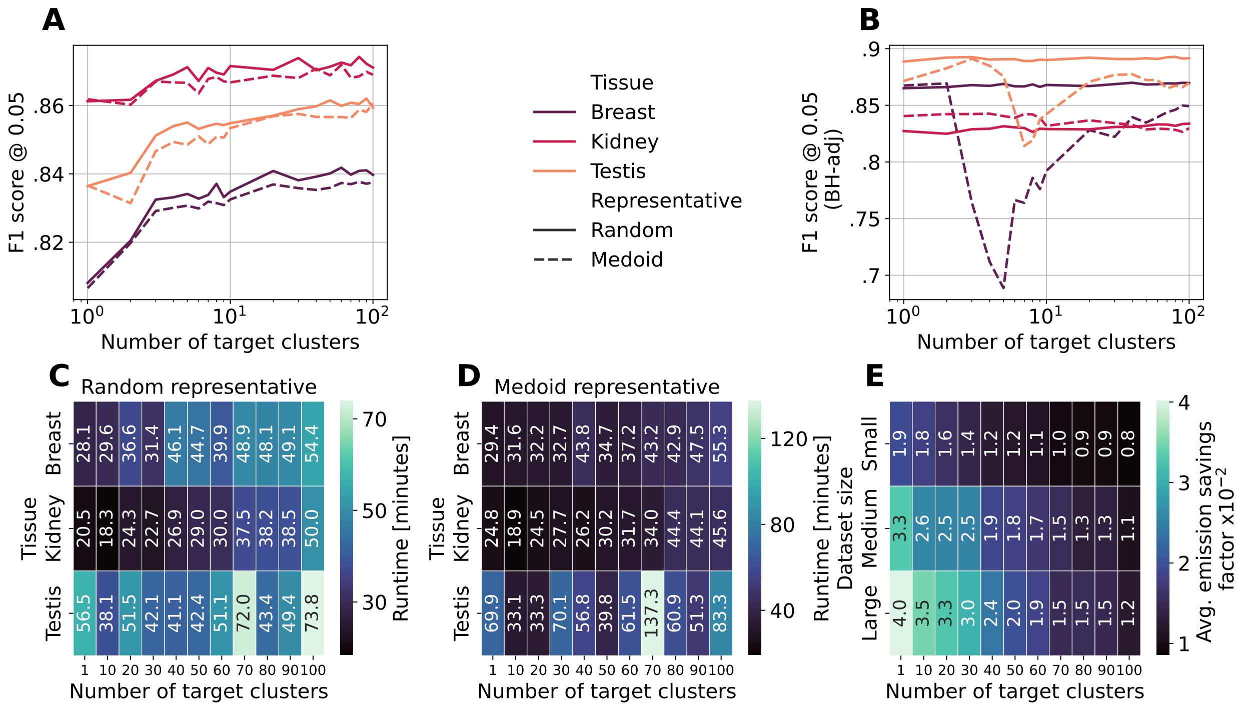


**Fig. B.1.** Additional results for SignifiKANTE with GRNBoost2 as core GRN inference method. (A, B, C, D) Effect of using cluster medoids or randomly sampled cluster genes as cluster representatives. (E) Saved carbon emission in comparison to $P$-value computation with DIANE-like permutation approach.

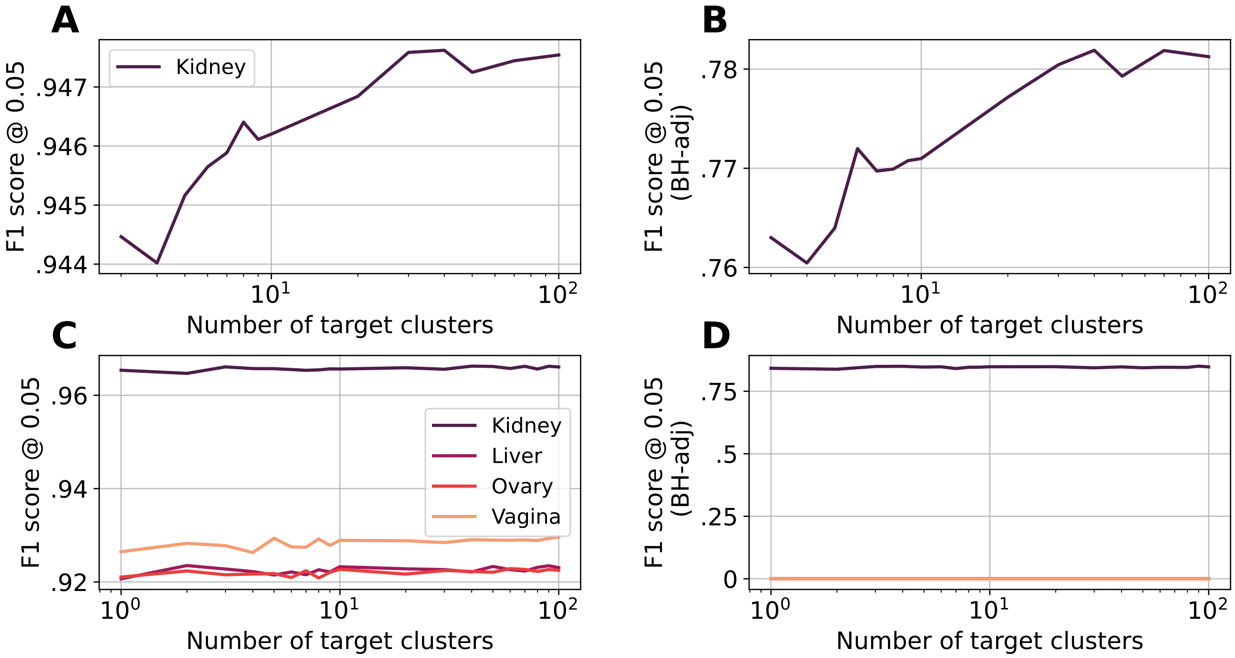


**Fig. B.2.** Results for SignifiKANTE with GENIE3 and GreNaDine's LASSO regression as core GRN inference method. (A, B) Results for GENIE3. Only the tissue kidney has been considered in this analysis, due to the immense overhead in runtime and memory consumption using GENIE3. For tissues with more samples, already the reference GRN could not be inferred below the job time limit of 24 hours of our compute cluster. The significant memory and runtime overhead of GENIE3 also led to SignifiKANTE not being able to successfully complete its $P$-value computation for a small number of $\ell$-values within the time frame of 24 hours. (C, D) Results for GreNaDine's LASSO regression. In (D), F1 scores for tissues liver, ovary, and vagina are zero due to the lack of significant edges (i.e. true positives) under Benjamini-Hochberg correction.

**Table B.1.** Mapping of GTEx [3] tissue names to terms in the BRENDA tissue ontology (BTO) [2]. The mapping was manually generated using semantic similarity between the terms.

| GTEx tissue name | BTO term | GTEx tissue name | BTO term |
|---|---|---|---|
| Adipose_Tissue | BTO:0001487 | Adrenal_Gland | BTO:0000047 |
| Bladder | BTO:0001418 | Blood | BTO:0000089 |
| Blood_Vessel | BTO:0001102 | Brain | BTO:0000142 |
| Breast | BTO:0000149 | Cervix_Uteri | BTO:0002249 |
| Colon | BTO:0000269 | Esophagus | BTO:0000959 |
| Fallopian_Tube | BTO:0000980 | Heart | BTO:0000562 |
| Kidney | BTO:0000671 | Liver | BTO:0000759 |
| Lung | BTO:0000763 | Muscle | BTO:0000887 |
| Nerve | BTO:0000925 | Ovary | BTO:0000975 |
| Pancreas | BTO:0000988 | Pituitary | BTO:0001073 |
| Prostate | BTO:0001129 | Salivary_Gland | BTO:0001203 |
| Skin | BTO:0001253 | Small_Intestine | BTO:0000651 |
| Spleen | BTO:0001281 | Stomach | BTO:0001307 |
| Testis | BTO:0001363 | Thyroid | BTO:0001379 |
| Uterus | BTO:0001424 | Vagina | BTO:0000243 |

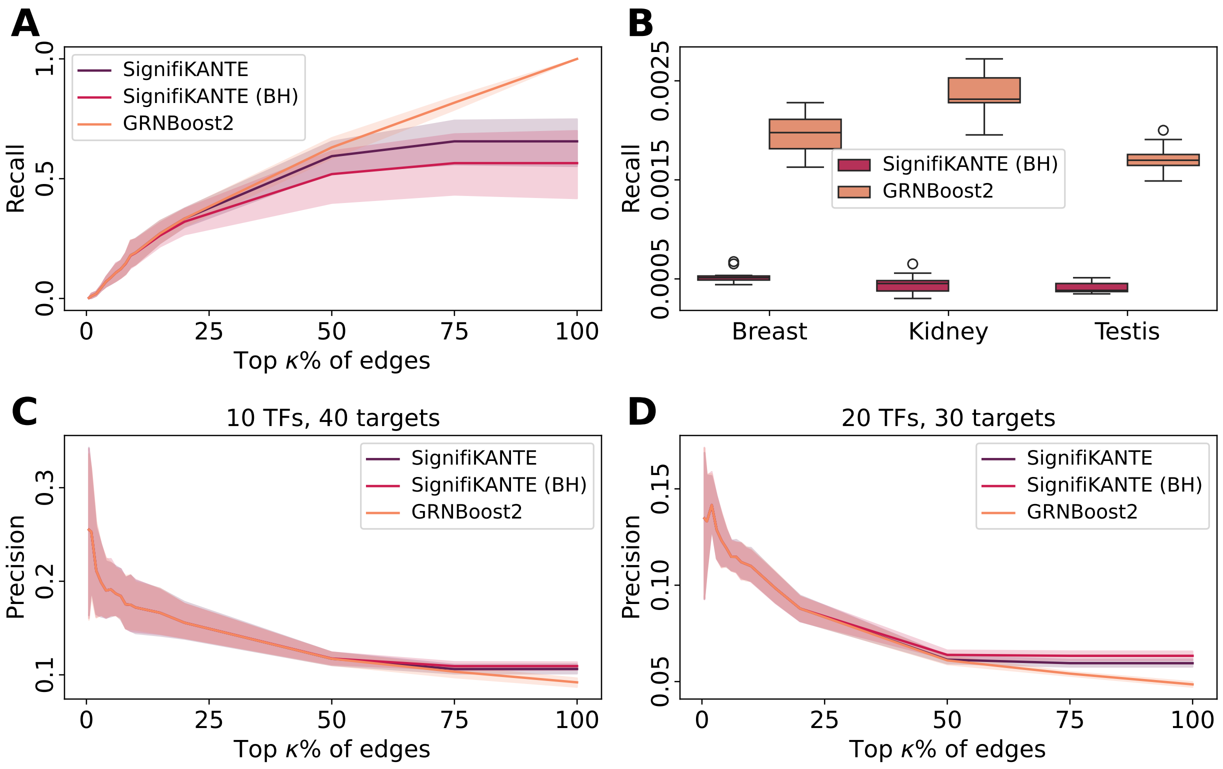


**Fig. B.3.** Additional results for simulated and real-world data. **(A)** Recall of top $\kappa\%$ edges (sorted by weights) inferred from simulated expression data against groundtruth GRN used for simulation, using all GRNBoost2 edges, all significant edges of SignifiKANTE (significance cutoff at $\alpha = 0.05$, $\ell = 10$ target gene clusters), and all significant edges of SignifiKANTE with BH correction (FDR control at $q = 0.05$, $\ell = 10$). **(B)** Distributions of recall values between all GRNBoost2 edges inferred from breast, kidney, and testis GTEx data and only significant edges computed by SignifiKANTE (BH correction, and FDR control at $q = 0.05$, $\ell = 100$). Known TF-target interactions from the CollecTRI database [19] were treated as groundtruth. Using different random seeds, we ran GRNBoost2 followed by SiginiKANTE ten times on the same data. **(C, D)** Additional precision values on simulated expression data, for the setting of 10 TFs with 40 targets each (C), and the setting of 20 TFs with 30 targets each (D).

**Table B.2.** Overview of GTEx data, including the number of samples and genes after preprocessing per tissue, and the number of edges and significant edges in the reference GRN computed by GRNBoost2. Significant edges are computed using SignifiKANTE's FDR control for significance level 0.05, $\ell = 100$, and Benjamini-Hochberg correction.

| Tissue | # Samples | # Genes | # Edges | # Significant edges |
|---|---|---|---|---|
| Adipose_Tissue | 1204 | 15620 | 1540796 | 340828 |
| Adrenal_Gland | 258 | 15630 | 1891813 | 307457 |
| Bladder | 21 | 15864 | 2304629 | 246183 |
| Blood | 929 | 13896 | 1724486 | 322745 |
| Blood_Vessel | 1335 | 15313 | 1501239 | 307109 |
| Brain | 2642 | 15884 | 2022744 | 608754 |
| Breast | 459 | 15755 | 1746068 | 285476 |
| Cervix_Uteri | 19 | 15672 | 2236599 | 190448 |
| Colon | 779 | 15888 | 1769981 | 354494 |
| Esophagus | 1445 | 15538 | 1765118 | 388435 |
| Fallopian_Tube | 9 | 15738 | 1947223 | 1063782 |
| Heart | 861 | 15216 | 1549753 | 446156 |
| Kidney | 89 | 16030 | 2079742 | 335425 |
| Liver | 226 | 15282 | 1818300 | 271637 |
| Lung | 578 | 16091 | 1751109 | 282877 |
| Muscle | 803 | 14852 | 1551872 | 282176 |
| Nerve | 619 | 15975 | 1694849 | 305257 |
| Ovary | 180 | 15599 | 2006770 | 205126 |
| Pancreas | 328 | 15606 | 1900876 | 420292 |
| Pituitary | 283 | 16448 | 1983108 | 287424 |
| Prostate | 245 | 16210 | 1977624 | 285565 |
| Salivary_Gland | 162 | 16123 | 2019478 | 287912 |
| Skin | 1809 | 15120 | 1994850 | 399343 |
| Small_Intestine | 187 | 16246 | 2067232 | 298896 |
| Spleen | 241 | 15945 | 1913360 | 315616 |
| Stomach | 359 | 15769 | 1902568 | 285085 |
| Testis | 361 | 18124 | 2049277 | 424118 |
| Thyroid | 653 | 15993 | 1711783 | 320693 |
| Uterus | 142 | 15681 | 2084537 | 179294 |
| Vagina | 156 | 15878 | 2135811 | 265521 |
